# Screening of functional maternal-specific chromatin regulators in early embryonic development

**DOI:** 10.1101/2024.03.06.583790

**Authors:** Guifen Liu, Yiman Wang, Xiangxiu Wang, Wen Wang, Zheng Cao, Yong Zhang

**Author notes:** Correspondence: Yong Zhang Ph.D. These authors contributed equally to this work (Guifen Liu, Yiman Wang, Xiangxiu Wang). Key Laboratory of Biorheological and Technology of Ministry of Education, State and Local Joint Engineering Laboratory for Vascular Implants, Modern Life Science Experiment Teaching Center at Bioengineering College of Chongqing University, Chongqing 400030, China.

## Abstract

The early stages of embryonic development rely on maternal products for proper regulation. However, systematic screening for functional maternal-specific factors has been challenging due to the time- and labor-intensive nature of traditional approaches. Here, we combined a computational pipeline and F0 homozygous mutation technology to screen for functional maternal-specific chromatin regulators in zebrafish embryogenesis and identified Mcm3l, Mcm6l, and Npm2a as playing essential roles in DNA replication and cell division. Our results contribute to understanding the molecular mechanisms underlying early embryo development and highlight the importance of maternal-specific chromatin regulators in this critical stage.

## Introduction

The genome is not yet active during the initial phase of embryonic development. Instead, maternal products (including RNAs and proteins) take charge of early development processes until zygotic genome activation (ZGA)^1,2^. Numerous studies have highlighted the importance of maternal products in the transition between maternal and embryonic control of development^1,3,4^. Among maternal products, a subset of genes show only maternal-specific expression^5,6^. A few maternal-specific genes have been identified and shown to significantly impact early embryonic development. For example, in mouse embryogenesis, maternal-specific factors Zar1^7^ and Npm2^8^ play roles in the 1-2 cell embryo, and Mater^9^, Zfp36l2^10^, and Oog1^11^ function in the 2-4 cell embryo, as well as Stella, which is involved in the morula development^12^. Several functional maternal-specific factors have been reported in zebrafish embryogenesis, including Foxr1^13^, Kpna7^14^, Slbp2^15^, Fue^16^, Plk1^17^, and Fhdc3^14^. While foundational works identified key maternal regulators of early embryogenesis^18,19^, subsequent studies built upon these findings further elucidating the molecular genetics of these maternal effects^20–23^. As additional maternal-specific factors may play indispensable roles in early embryogenesis, a systematic screening to identify functional maternal-specific genes is in demand.

Traditional approaches to studying the function of maternal-specific factors require multi-generation hybridization screening to obtain homozygous gene mutations, which are time-consuming and labor-intensive. Recently, the CRISPR/Cas9 system has been successfully applied in several species to generate homozygous mutations in F0 embryos^24,25^, which is ideal for systematic screening of functional maternal-specific factors in early development. Zebrafish is widely used for developmental studies because it allows rapid and efficient *in vitro* manipulations. As zebrafish embryos undergo rapid cleavage^26^, maternal-specific chromatin regulators may be functionally essential for maintaining genome integrity and proper genome activation. In this study, we combined F0 homozygous mutation technology with a computational pipeline to screen for the functional maternal-specific chromatin regulators in zebrafish embryogenesis, and we identified the maternal-specific chromatin regulators Mcm3l, Mcm6l, and Npm2a as critical for early zebrafish embryonic development.

## Results

### Screening for maternal-specific chromatin regulators

To systematically screen for maternal-specific chromatin regulators, we first collected publicly available gene expression datasets in multiple developmental stages and adult tissues of zebrafish (Table S1). The datasets were divided into the early embryo group and the late embryo & adult tissues group. The early embryo group included datasets from MII oocytes to before 24 hpf embryos. In comparison, the late embryo & adult tissues group had datasets from 24 hpf embryos to adult tissues, including heart, muscle, liver, brain, intestinal epithelium, spinal cord, gills, caudal fin, and blood. We then calculated the expression levels of all annotated coding exons for all datasets (see Methods for details) and defined potential maternal-specific exons as those with high expression levels (TPM > 6.0) in at least one dataset in the early embryo group and low expression levels (TPM < 0.8) in all datasets in the late embryo & adult tissues group (Fig 1A). 361 genes with at least one potential maternal-specific exon were identified (Fig S1A, Table S2). Since some identified genes may also be zygotically transcribed genes, we applied two approaches to filter out the zygotically transcribed genes (Fig 1A). First, because H3K36me3 and RNA Pol II are associated with transcription elongation^27,28^, we collected their ChIP-seq data at dome stage^29^ and filtered out genes with strong H3K36me3 or RNA Pol II signals (Fig 1A, S1B, Table S1, S2; see Methods for details). Second, we collected lists of zygotically transcribed genes from the literature^30^ and filtered out genes reported as zygotically transcribed genes (Fig 1A). After filtering, 178 genes remained as candidate maternal-specific genes (Table S2).

**Figure 1.**
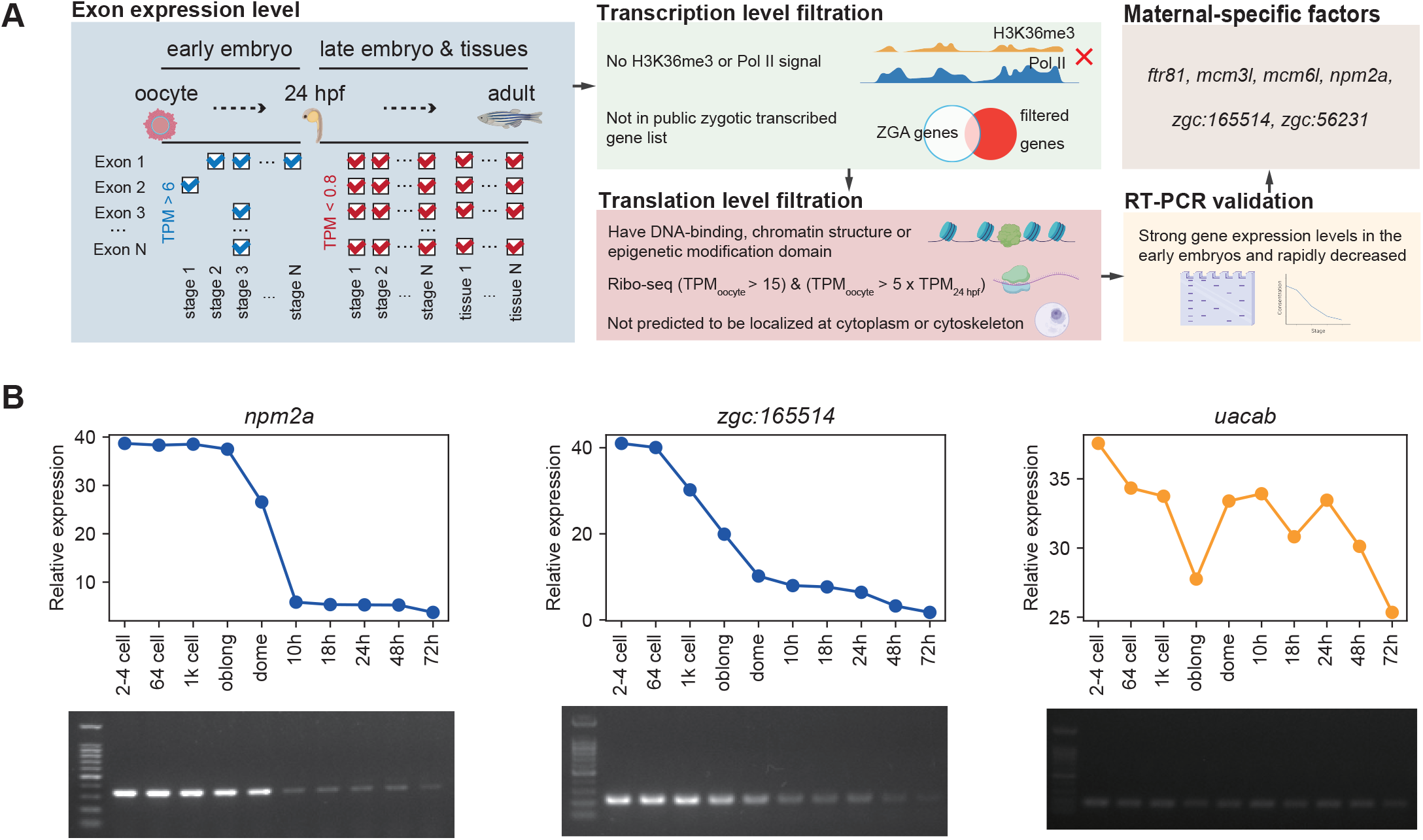
Screening for maternal-specific chromatin regulators. (*A*) Schematic of maternal-specific chromatin regulator screening. (*B*) Gel electrophoresis image (top) and quantitative statistics of the fluorescence intensity by ImageJ (bottom) of examples of candidate maternal-specific chromatin regulator expression by RT-PCR at 2-4 cell, 64-cell, 1k cell, oblong, dome, 10 hpf, 18 hpf, 24 hpf, 48 hpf, and 3 dpf. The expression levels of *npm2a* and *zgc:165514* continuously decrease during early development.

Since chromatin regulators, including transcription factors and epigenetic modifiers, play essential roles in early zebrafish embryo development^31,32^, we focused on maternal-specific chromatin regulators in this study. For each candidate maternal-specific gene, we used the NCBI Conserved Domain Database (CDD)^33^ to examine whether its protein sequence contained a DNA-binding domain or a domain related to chromatin structure or modification (Fig 1A), and 30 potential maternal-specific chromatin regulators were identified (Fig S1C; Table S2). We further investigated the translational activities of the potential maternal-specific chromatin regulators by re-analyzing Ribo-seq datasets during zebrafish embryonic development^34,35^ (see Methods for details). We set a threshold based on the Ribo-seq datasets (TPM_0 hpf_ > 15 & TPM_0 hpf_ > 5 x TPM_24 hpf_) for chromatin regulators with high maternal translational activities, and 21 potential maternal-specific chromatin regulators were selected (Fig 1A, S1D, Table S2; see Methods for details). Next, we excluded factors whose predicted subcellular localizations are only cytoplasmic or cytoskeletal according to the annotations in ZFIN^36^ (Fig 1A, Table S2), and 15 potential maternal-specific chromatin regulators were retained.

We further validated the expression levels of the genes of potential maternal-specific chromatin regulators by RT-PCR in 2-4 cell, 64-cell, 1k cell, oblong, dome, 10 hpf, 18 hpf, 24hpf, 48 hpf, and 3 dpf embryos (Fig 1B, S1E). RT-PCR analysis confirmed that 6 chromatin regulators, *i*.*e*., Ftr81, Mcm3l, Mcm6l, Npm2a, Zgc:165514, and Zgc:56231, exhibited intense gene expression levels in the early embryos, while their gene expression levels rapidly decreased to undetectable levels after 24 hpf. These 6 maternal-specific chromatin regulators were selected for functional evaluation.

### Functional evaluation of maternal-specific chromatin regulators

Since the technology to generate homozygous mutations in F0 generation embryos is ideal for systematic screening of functional maternal-specific factors in early development^24,25^, we first validated its effectiveness in disrupting the pigment gene *tyr* in zebrafish. We designed 6 adjacent sgRNAs on exon 1 and injected Cas9 with single sgRNA or different combinations of sgRNAs, respectively. The mutation efficiency can reach 100% when using a combination of 4 sgRNAs (Table S3; see Methods for details). It should be noted that co-injection of 4 adjacent sgRNAs resulted in large deletions, with 97.1% of mutants showing deletions larger than 20 bp, 52.9% showing deletions larger than 50 bp, and 23.5% showing deletions larger than 100 bp. In contrast, the deletion sizes were usually smaller than 10 bp when a single sgRNA was injected (Fig S2A, B). Taking advantage of the large deletion sizes induced by co-injected sgRNAs, we accelerated the detection of mutation efficiency by observing the dispersed DNA fragments by high-concentration gel electrophoresis (Fig S2C; see Methods for details), which can avoid the traditional costly and time-consuming TA cloning sequencing. We then tested the feasibility of this mutation efficiency detection approach for 2 genes with known knockout phenotypes, *smarca4a* (cardiac edema, short body) and *ddx19* (slender tail, outward curvature). The consistency between phenotype and mutation efficiency detection confirmed that this approach is generally applicable for mutation efficiency detection (Fig S2D, E). Our results showed we could successfully and efficiently generate homozygous mutations in F0 embryos.

Based on the validated method to generate homozygous mutations in F0 embryos, we performed homozygous knockout for the genes of 6 maternal-specific chromatin regulators (Fig 2A, B, Fig S2F). These 6 F0 mutant lines, more than 50 for each line, were cultured under standard laboratory conditions. Notably, none of the 6 F0 mutant lines exhibited any obvious phenotype and were successfully raised to sexual maturity. The mutant females were mated with wild-type males to generate F1 generation embryos with corresponding maternal deletions. Phenotypic observation revealed that the F1 embryos with maternal deletion of three genes (*mcm3l, mcm6l*, and *npm2a*) arrested their development around the mid-blastula transition (MBT) and failed to undergo the entire ZGA process, resulting in death at 7.5 hpf (Fig 2C, Fig S2G). Furthermore, the expression level of several maternal genes and zygotic genes in *mcm3l, mcm6l* and *npm2a* mutant embryos also demonstrated a significant delay or disruption in maternal transcript degradation, coupled with considerable weakening of zygotic transcript expression in these mutants (Fig S2H, I). Taken together, we evaluated the functions of maternal-specific chromatin regulators and demonstrated that 3 have essential effects on early embryonic development.

**Figure 2.**
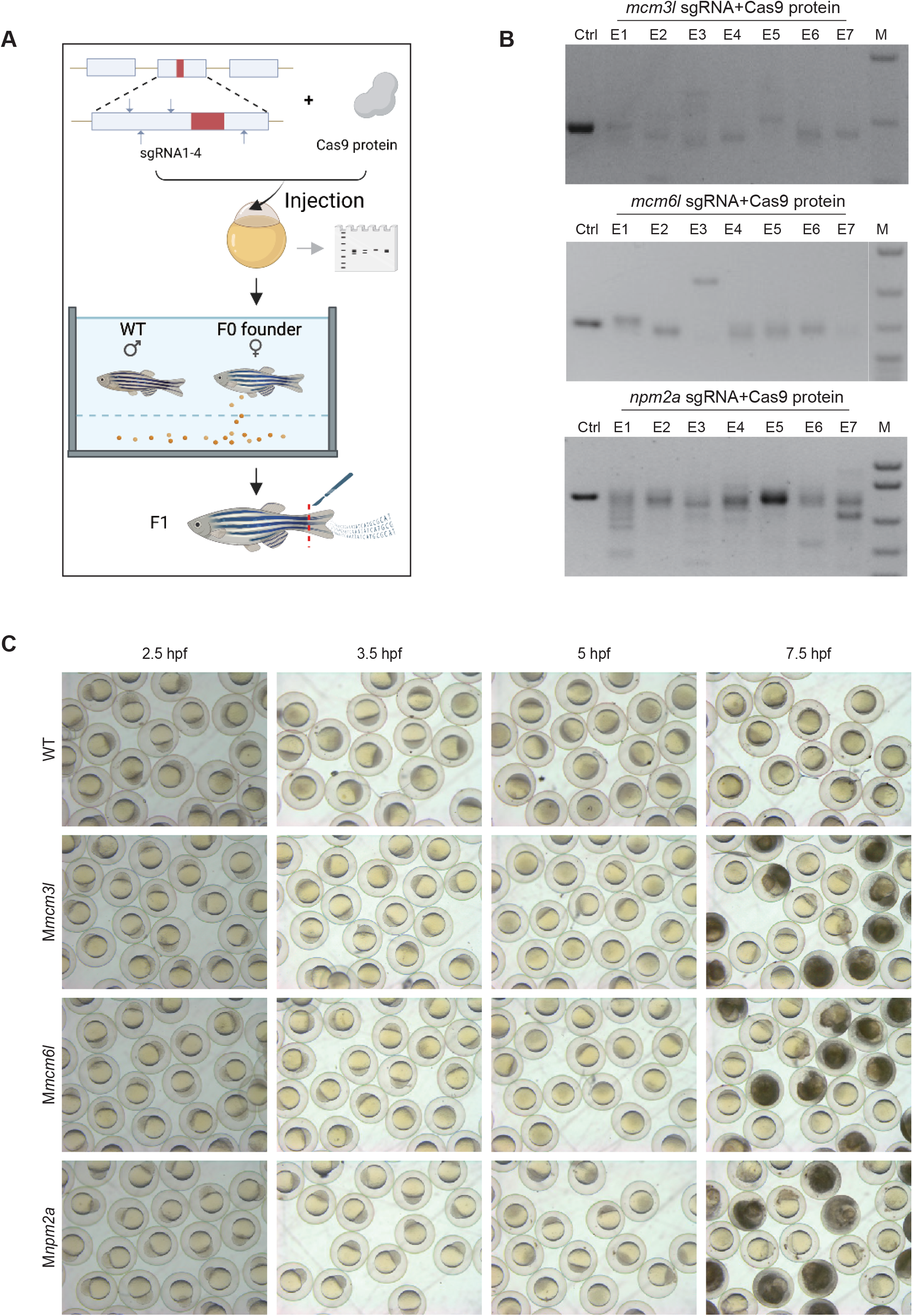
Functional evaluation of maternal-specific chromatin regulators. (*A*) Schematic of construction for F0 homozygous mutants using CRISPR/Cas9. (*B*) Gel electrophoresis image of examples of genotypic mutation efficiency using a single embryo for *npm2a, mcm3l*, and *mcm6l* at 24 hpf. (*C*) Photos of developmental morphology for F1 generation embryos for the WT and maternal-specific chromatin regulator homozygous mutants, including *mcm3l* (top), *mcm6l* (middle) and *npm2a* (bottom).

### The role of Mcm6l in DNA replication in early embryos

Among the 3 functional maternal-specific chromatin regulators, Mcm3l and Mcm6l are the homologous proteins of Mcm3 and Mcm6 in the mini-chromosome maintenance (MCM) family. Since the expressions of *mcm3* and *mcm6* are undetectable before ZGA^37^, we hypothesized that Mcm3l and Mcm6l might play a critical role related to the rapid DNA replication before ZGA. A recent study showed that nuclear division is disrupted in the *mcm3l* mutant, and most cells undergo multiple anucleate divisions^17^. We next investigated whether *mcm6l* mutation can affect DNA replication before ZGA. We performed a relative quantitative analysis of DNA content by whole genome gel electrophoresis, and conducted the qRT-PCR to assess the expression level of *pcna* and histone 3. The observed increase in the expression levels of *pcna* and *h3* over time in wild-type specimens, but not in M*mcm6l* mutants, demonstrated that DNA replication stopped at 1.5 hpf in *mcm6l* mutant embryos (Fig 3A, B), indicating that Mcm6l can delay or arrest DNA replication at the early stages of embryogenesis. We then detected the nuclear division of embryos at different stages by DAPI staining. We found that the nuclei of *mcm6l* mutant embryos were abnormal from the 1-cell stage (0 hpf), showing diffuse granules (Fig 3C). In *mcm6l* mutant embryos, aggregated nuclei were difficult to find, with only small DAPI-positive fragments visible at 2 hpf, and then DAPI-negative, larger-than-normal structures appeared at 4.5 hpf (Fig 3B). Further cell membrane staining with phalloidin revealed an intact cell membrane structure without a typical nucleus for the *mcm6l* mutant embryo (Fig S3). Our results suggested that, similar to *mcm3l*, deletion of *mcm6l* can cause DNA replication arrest and cell anucleation, ultimately leading to embryonic developmental arrest at MBT.

**Figure 3.**
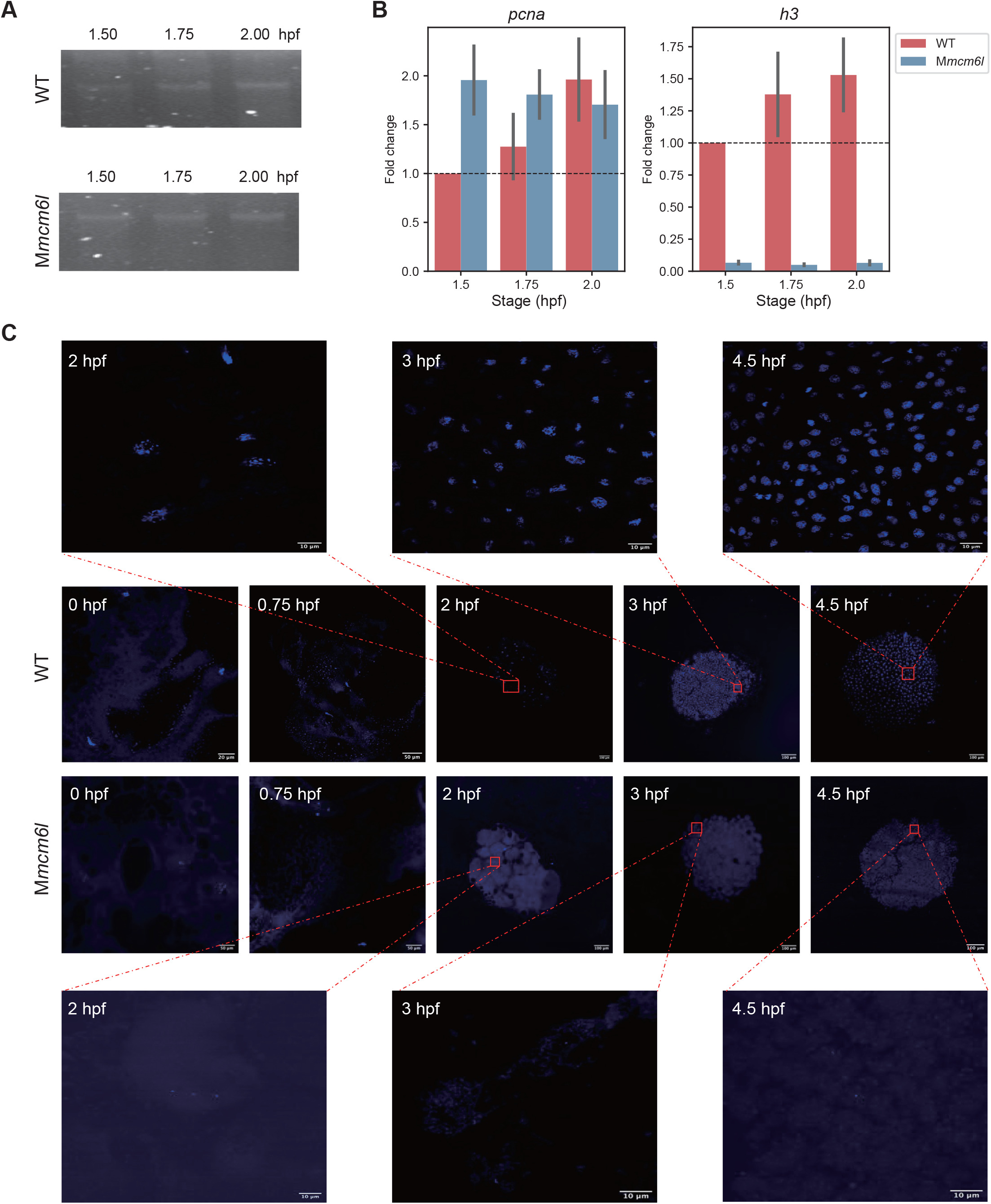
The role of Mcm6l in DNA replication in early embryos. (*A*) Gel electrophoresis of genomic DNA detection for WT and *mcm6l* F1 embryos at different developmental stages (1.5 hpf, 1.75 hpf, and 2.00 hpf). (*B*) The results of RT-qPCR for *pcna* and *h3* at 1.5 hpf, 1.75 hpf and 2.00 hpf. Comparing with WT, the expression level for *pcna* and *h3* did not exhibit an increasing trend during early development. (*C*) Nuclear staining of WT and M*mcm6l* F1 embryos at 0 hpf, 0.75 hpf, 2 hpf, 3 hpf and 4.5 hpf developmental stages. High magnified views of image regions marked by rectangles at different developmental stages of *mcm6l* mutant are shown.

### The role of Npm2a in cell division in early embryos

Another functional maternal-specific chromatin regulator, Npm2a, is a member of the nucleoplasmin family. A recent study showed that the deletion of *npm2a* resulted in reduced oocyte competence, with most spawned oocytes failing to divide^38^, which is inconsistent with our observations that *npm2a* mutant embryos underwent several divisions and arrested their development around the MBT (Fig 2C). The nuclei of *npm2a* mutant embryos appeared normal at the 1-cell stage (0 hpf) and 0.75 hpf. At the 2 hpf, the nuclei showed many disordered, filamentous, and partially aggregated DNA. However, by 3 hpf, the nuclei tended to become more intact, and the filamentous structures were gradually reduced. A group of DAPI-negative, larger-than-normal structures appeared at 4.5 hpf, along with numerous nuclear structures that were morphologically normal but different in size. The distribution of these two distinct types of structures was apparent, as they were found on separate sides of the embryo rather than intermingled (Fig 4A). Cell membrane staining with phalloidin revealed three types of membrane and nuclear structures: partially intact cell membrane and nuclear structure, partially intact cell membrane structure without a typical nucleus, and partially disordered filamentous nucleus-like structure that could not be merged with cells (Fig S4A). We further examined the DNA content and observed the delay in DNA replication in *npm2a* mutant embryos, whose DNA content at 5.5 hpf was similar to that at the 1k-cell stage (3 hpf) in wildtype (WT) embryos (Fig 4B). We next performed RNA-seq in *npm2a* mutant and WT embryos at 3.66 hpf (*i*.*e*., oblong stage for WT embryos) to examine whether the transcriptional activations of these embryos are affected. To exclude the influence of maternal-loading mRNAs, we focused on genes only expressed zygotically (referred to as zygotic genes). Compared to genes expressed both maternally and zygotically^30^ (referred to as maternal-zygotic genes), zygotic genes tended to decrease their expression levels in *npm2a* mutant embryos (Fig S4B). Our results demonstrated that deletion of *npm2a* can lead to DNA replication delay and abnormal nuclear division, resulting in the arrest of embryonic development around the MBT.

**Figure 4.**
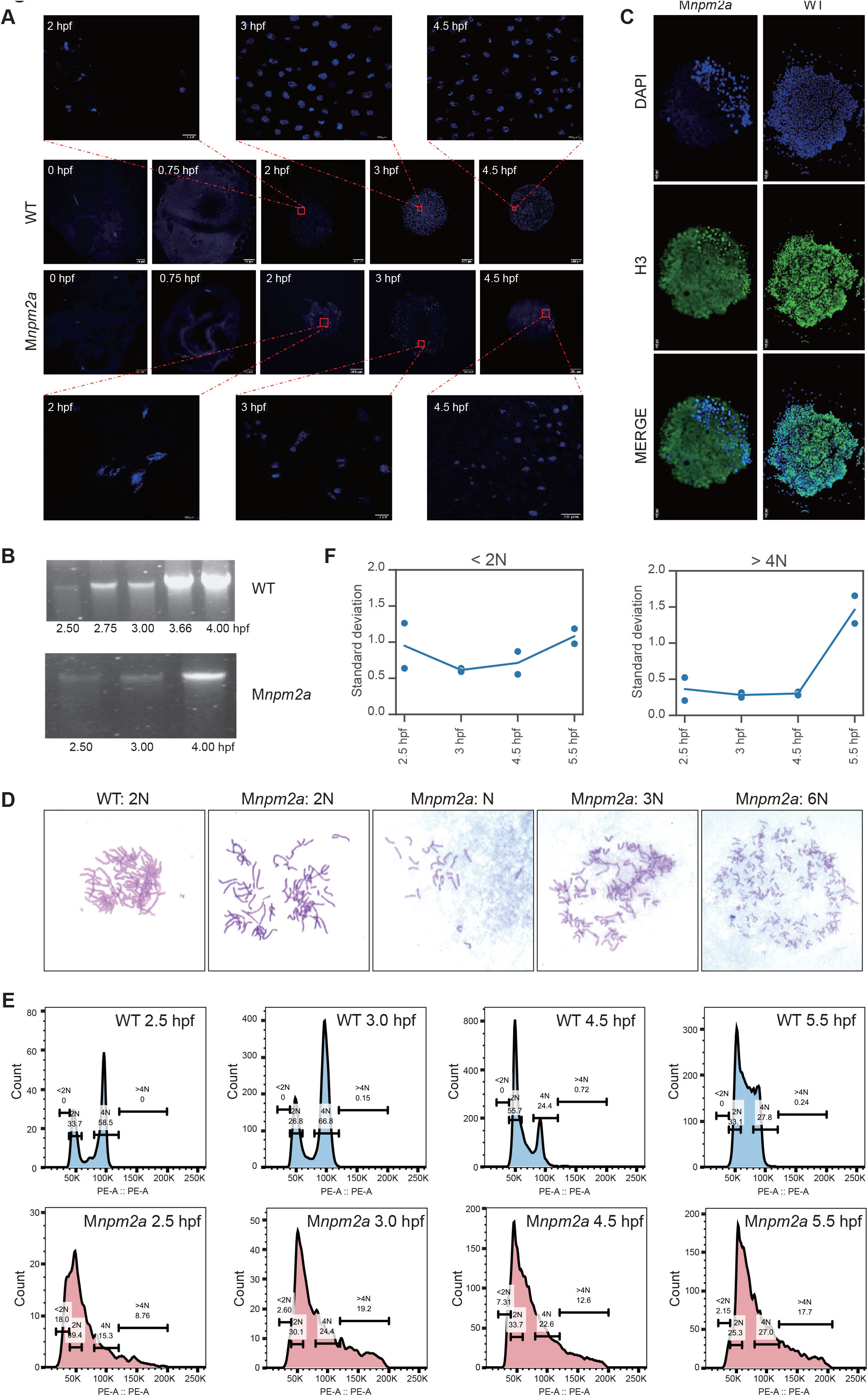
The role of Npm2a in cell division in early embryos. (*A*) Nuclear staining of WT and M*npm2a* F1 embryos at developmental stages of 0 hpf, 0.75 hpf, 2 hpf, 3 hpf and 4.5 hpf. High magnified views of image regions marked by rectangles at different developmental stages of *npm2a* mutant are shown. (*B*) Gel electrophoresis of genomic DNA detection for WT and *npm2a* F1 embryos at different developmental stages (2.50 hpf, 2.75 hpf, 3.00 hpf, 3.66 hpf and 4.00 hpf for WT; 2.50 hpf, 3.00 hpf and 4 hpf for *npm2a* mutants). (*C*) The immunofluorescence staining of h3 (green) for *npm2a* mutants (left) and WT (right) at 4.5 hpf. The nuclei were stained with DAPI (blue). (*D*) Representative images showing karyotype analysis of *npm2a* mutant embryos. n=25 (*E*) DNA content analysis by flow cytometry (PI staining) of WT (top) and M*npm2a* F1 embryos (bottom) at 2.5 hpf, 3.0 hpf, 4.5 hpf, and 5.5 hpf. The channel of PI is PE-A. Excitation: 488 nm. Emission: 564 nm. (*F*) Standard deviation of the log_2_ fold-change of the observed coverage and expected coverage for each chromosome of cells with less than 2N DNA content (left) and more than 4N DNA content (right).

To further investigate the abnormal nuclear division in *npm2a* mutant embryos, we first examined the state for chromosomes through H3 immunofluorescence. This analysis revealed that DAPI-positive cells also exhibited H3 fluorescence, while DAPI-negative cells lacked H3 fluorescence, indicating partial visualization of chromosomes (Fig 4C). Additionally, karyotype analysis revealed abnormal chromosomes, including hexaploidy (6N), triploid (3N) and haploid (N) in certain cells (Fig 4D). Then we examined the distribution of DNA contents in cells by using PI staining followed by flow cytometry at four developmental stages (2.5 hpf, 3 hpf, 4.5 hpf, and 5.5 hpf). We observed that a significant population of cells from *npm2a* mutant embryos have a DNA content greater than 4N or less than 2N (Fig 4E). To examine whether the abnormal nuclear division in *npm2a* mutant embryos has a chromatin preference, we performed whole genome sequencing (WGS) for cells with DNA content greater than 4N or less than 2N for each developmental stage (Table S4). We calculated the ratio of observed and expected read counts for each chromosome and found no consistent patterns between biological replicates, indicating random chromatin distribution during abnormal nuclear division (Fig S4C; see Methods for details). However, we observed an accumulation of chromatin distribution imbalance around the MBT, with samples at 5.5 hpf exhibiting the highest variation in chromatin distribution (Fig 4F). To investigate whether the deletion of *npm2a* leads to defects in genome integrity, we further calculated the differences in observed-to-expected read count ratios between adjacent 1-Mb genomic windows. We observed a significant proportion of ratio differences larger than 2-fold, particularly in greater than 4N samples at 5.5 hpf and most less than 2N samples, indicating widespread truncated chromatin in those samples (Fig S4D; see Methods for details). Taken together, our results suggest that Npm2a plays an essential role in chromosome segregation and genome integrity during early embryogenesis.

## Discussion

This study introduced a systematic approach to identify and functionally evaluate maternal-specific chromatin regulators in zebrafish embryogenesis, leading to the discovery of Mcm3l, Mcm6l, and Npm2a as critical factors in early embryo development. By combining computational screening with F0 homozygous mutation technology, this time- and labor-efficient approach is particularly suited for uncovering the functions of maternal-specific factors. Chromatin regulators represent only a small percentage of candidate maternal-specific factors, and there are over 100 non-chromatin-regulator factors were identified and listed in Table S2, allowing researchers to further explore their functions. Furthermore, the approach presented in this study could be effectively applied to uncover the functions of maternal-specific factors in other organisms.

The early embryos of zebrafish undergo rapid cellular cleavage without cell cycle checkpoints until the MBT^39^, and studies have shown the existence of asynthetic fission in early embryos, where cells divide with minimal or no DNA synthesis^16,17,40^. In this study, we found that the deletion of *mcm6l* and *npm2a* can affect DNA replication or nuclear division, leading to many anucleated cells. Mcm3l, the cleavage-stage-specific counterpart of Mcm3, is known to be essential for the initiation of DNA replication^17^. Given the similarity in phenotype observed between *mcm6l* and *mcm3l* mutant embryos, we hypothesized that Mcm6l may also be important for rapid DNA replication and lack of cell cycle checkpoints before MBT. In contrast to a recent study^38^, we observed that the deletion of *npm2a* led to delayed DNA replication and abnormal nuclear division, ultimately resulting in embryonic development arrest at the MBT. Potential reasons for discrepancies in phenotypes include variability in CRISPR/Cas9 editing efficiency and specificity, as well as the possibility that *npm2a* functions redundantly with other genes or pathways. Since nucleoplasmin is required for mitotic progression^41^, Npm2a likely plays a critical role in chromosome segregation and genome integrity during rapid cellular cleavage that occurs in early embryos. However, further investigation is needed to determine the mechanisms that lead to abnormal nuclear division induced by *npm2a* mutation in early embryos.

## Method

### Zebrafish husbandry

Wild-type Tübingen-strain zebrafish were maintained, raised, and crossed under standard condition^42^. Zebrafish embryos were collected at the one-stage and allowed to develop to the desired stage at 28.5°C. Zebrafish care and experiments were approved by the Institutional Animal Care and Use Committee of Tongji University.

### RNA extraction, PCR and RNA-seq library preparation and sequencing

Embryos developed to desired stages were collected and removed from their chorions (50 embryos for each stage). RNA was lysed by thorough homogenization in 500 *μ*l RNAiso Plus (TAKARA 9108), extracted with chloroform, precipitated by isopropyl alcohol, followed by washing with 75% ethanol, and dissolved in nuclease-free water (Invitrogen AM9930). The cDNA library was synthesized using EvoScript Universal cDNA Master (Roche 07912374001). PCR were performed using 2x EasyTaq PCR SuperMix (TransGen AS111), and detected by agarose gel electrophoresis. qPCR was conducted using SYBR Green Master (Roche 04913914001) and were measured by 2^(−ΔΔCt))^. All primer sequences for RT-PCR are listed in Table S5.

The RNA-seq libraries of zebrafish embryos were prepared according to the protocols for the KAPA mRNA HyperPrep Kit (KAPA Biosystems KK8580). The amplified libraries were sequenced by the Illumina NovaSeq platform.

### CRISPR/Cas9-mediated mutation, Single-cell PCR Analysis, TA Cloning and Genotyping

The guide RNAs sequences were designed according to a previously study^25^, synthesized using 2x Pfu Master Mix (novoprotein E006), and subjected to in vitro transcription with the MEGAshortscript T7 kit (Invitrogen AM1354). Cas9 protein (NEB, 0646T) (4 μM) and sgRNA (each 60 ng/μl) were injected into zebrafish embryos at the one-cell stage. Single 24 hpf embryos or adult zebrafish tails were picked up in a PCR tube, and the embryos were lysed using 20 μl lysis buffer (100 mM Tris-HCl, pH 8.3; 200 mM NaCl; 0.4% SDS; 5 mM EDTA) containing proteinase K at 56°C for 4h or overnight, followed by heat inactivation of proteinase K at 95°C for 15 min. The lysis products were then used as templates for subsequent PCR analysis with 2x EasyTaq PCR SuperMix (TransGen AS111). The PCR products were analyzed by 2% agarose gel electrophoresis for mutation efficiency detection, or the gel can be purified and cloned using pEASY-T5 Zero Cloning Kit (TransGen CT501) following the manufacturer’s instructions. Colonies were picked from each transformation and Sanger sequencing was applied to detect genotype. All sgRNA and primer sequences are listed in Table S5.

### Immunofluorescence, nuclear staining and cell membrane staining

WT or mutant embryos were collected at desired stages, removed from their chorions by pronase (1mg/ml) and deyolked manually. The cell mass was added onto the polylysine-coated coverslips and allowed to settle for 5-10min at 50°C. Subsequently, the samples were fixed with 4% paraformaldehyde for 20 min at room temperature, and washed in PBS three times, 5min for each wash. Then, the samples were permeabilized and blocked with a solution containing 5% goat serum, 0.1% Triton® X-100, They were then incubated with an H3 antibody (Cell Signaling #4499), followed by a secondary antibody. Cell membrane staining was performed with phalloidin (Invitrogen™ A12379) following the manufacturer’s instructions. Finally, the nuclear DNA were stained with DAPI and imaged using a microscope (Olympus BX53).

### Genome DNA extraction

The embryos at the desired stage were collected and lysed by gDNA lysis buffer (100 mM Tris-HCl, pH 8.3; 200 mM NaCl; 0.4% SDS; 5 mM EDTA), containing proteinase K at 56°C for 4h or overnight. Then the gDNA was extracted by Phenol/Chloroform/Isoamyl alcohol (25:24:1), and detected by electrophoresis.

### Karyotype analysis

The npm2a mutant embryos were dechorionated using pronase (1mg/ml) at 2.0 hpf. Add colchicine to a final concentration of 0.5 μM and incubate for 60 min. Then we obtained a ball of cells by breaking the yolks through lifting the embryos to the surface. Subsequently, a hypotonic solution (75 mM KCl) was added to the cells and incubated at room temperature for 30 min. Fix the cells in a 3:1 methanol:acetic acid and stained with Giemsa.

### DNA content analyses by flow cytometry, sorting and genome-seq

The embryos were collected at the desired stages, removed from their chorions and deyolked manually. The cell mass was picked up into a 1.5 ml centrifugal tube, and gently suspended with FxCycle PI/RNase solution (Invitrogen F10797). Then the DNA content was detected by flow cytometry (BD FACSVerse), and the data were analyzed using FlowJo software. For target cells, flow cytometry sorting was carried out (BD FACSAria III), and the whole genome was amplified through REPLI-g Single Cell Kit (Qiagen 150343). Then the genome DNA were fragmented by covaries (M220), and the library was constructed using HyperPrep Kit (KAPA Biosystems KK8504) and sequenced by the Illumina NovaSeq platform.

### Processing of RNA-seq data

The RNA-seq data (Table S1) were subjected to filtration using TrimGalore (version 0.6.5, http://singlecellqc.com/projects/trim_galore/) along with cutadapt (version 3.7)^43^, applying the parameters “--trim-n”. Using HISAT2 (version 2.1.0)^44^ and applying the parameters “--no-mixed --no-discordant”, the filtered reads were remapped onto the zebrafish genome (danRer11 with an updated assembly of Chromosome 4^45,46^). StringTie (version 1.3.3b)^47,48^ was utilized to determine the TPM for each exon. The ComBat-seq^49^ method from Bioconductor sva^50^ (v3.36.0) was used to remove the batch effect for datasets from the same stage and tissue and replicates that had a low correlation (<0.8) with other replicates from the same stage or tissue were removed. The expression level of exons was averaged by replicates from the same stage or tissue. The processed Ribo-seq data was obtained from https://www.giraldezlab.org/data/ribosome_profiling/ ^35^. Data of 0 hpf and 24 hpf were used in this study.

### Processing of ChIP-seq data

Both H3K36me3 and RNA Pol II ChIP-seq data at dome stage (Table S1) were filtered by TrimGalore (version 0.6.5, http://singlecellqc.com/projects/trim_galore/) with cutadapt (version 3.7)^43^ with the following parameters: “--trim-n”. Bowtie 2 (version 2.4.2)^51^ was utilized to map the filtered reads back to the genome (danRer11 with new assembly Chromosome 4 for zebrafish data^45,46^) using the parameters “--no-mixed -- no-discordant --no-unal”. Reads with MAPQ scores less than 30 were excluded and SAMtools (version 1.6)^52,53^ was utilized to convert them to BAM format. Utilizing BEDTools (version 2.27.1)^54^, BED files were generated by converting the BAM files. Estimated fragment lengths were obtained from the log file of MACS2 (version 2.1.1.20160309)^55^ using the parameter “-f BED -g 1.34e9 -q 0.05”. The estimated fragment length was included in the reads, and the middle 50% was stacked and converted to bigWig for easy viewing and further examination through customized scripts and BEDTools (version 2.27.1)^54^. Maternal-specific exons with RPKM of both H3K36me3 in the last 2/3 regions of concatenated exons and RNA Pol II in the promoter regions lower than 0.2 at the dome stage were selected.

### Processing of Whole-Genome Sequencing (WGS) data

The reads of WGS data were filtered by TrimGalore (version 0.6.5, http://singlecellqc.com/projects/trim_galore/) with cutadapt (version 3.7)^43^ using the following parameters: “--trim-n”. Bowtie 2 (version 2.4.2)^51^ was utilized to map the filtered reads back to the genome (danRer11 with new assembly Chromosome 4 for zebrafish data^45,46^) using default parameters. SAMtools (version 1.6)^52,53^ were used to convert the reads to BAM format. Utilizing BEDTools genomecov module (version 2.27.1)^54^, BedGraph files were generated by converting the BAM files. The observed coverage of chromosomes or 1 Mb bins was calculated using the BedGraph files. The expected coverage was averaged using observed coverage and weighted by chromosome lengths or bin length.

### UCSC genome browser

The genome browser view was obtained using the UCSC genome browser^56,57^ with Track data hubs^58^, and visualized by applying a smoothing with a mean of pixels.

### Statistical analysis

*P* values were calculated by two-sided Mann-Whitney rank test using SciPy (version 1.5.4)^59^. Asterisk presents statistical significance (****: p-value < 0.0001; ***: p-value < 0.001; **: p-value < 0.01; *: p-value < 0.05; n.s.: not significant).

### Raw sequencing data access

The raw sequence data reported in this paper have been deposited in the Genome Sequence Archive^60^ in National Genomics Data Center^61^, China National Center for Bioinformation / Beijing Institute of Genomics, Chinese Academy of Sciences (GSA: CRA010494, shared URL for reviewers is https://ngdc.cncb.ac.cn/gsa/s/42EcBcB9, valid before July-3-2023) that are publicly accessible at https://ngdc.cncb.ac.cn/gsa.

## Acknowledgments

We thank the advice from Youwei Chen. This work was supported by the National Natural Science Foundation of China (32030022, 31970642) and the National Key Research and Development Program of China (2021YFA1302500).

## Author contributions

Y.Z. conceived and designed the study. G.L and X.W performed the experiments, with the help of Z.C. The data were analyzed by Y.W and W.W. The manuscript was written by G.L., Y.W., and Y.Z.

## Supplementary tables

**Table S1**. Public datasets used for maternal-specific chromatin-regulator screening.

**Table S2**. Results of filtration steps of maternal-specific chromatin regulators.

**Table S3**. Comparison of knockout efficiency between single gRNA and multiple gRNAs.

**Table S4**. High-throughput sequencing data generated in this study.

**Table S5**. Sequence of PCR primers and sgRNAs.

